# Pervasive suppressors halt the spread of selfish *Segregation Distorter* in a natural population

**DOI:** 10.1101/2025.10.13.681989

**Authors:** Ching-Ho Chang, Tyler Handler, Nick Fuda, Danielle Pascua, Taylar Mouton, Amanda M Larracuente

## Abstract

Meiotic drivers are selfish genetic elements that subvert Mendelian inheritance to increase their own transmission, yet they are typically found at low frequencies across natural populations. The factors that limit their spread remain unclear. To investigate this paradox, we studied the *Segregation Distorter (SD)* system, a selfish coadapted gene complex in *Drosophila melanogaster. SD* biases its transmission by killing sperm carrying a homologous chromosome bearing a target locus, *Responder* (*Rsp*), which appear as satellite repeats. Such selfish killing impairs male fertility and imposes selective pressure on the host genome to evolve resistance, either by deleting *Rsp* copies or acquiring unlinked suppressors. To characterize the spectrum of *Rsp* alleles and the frequency of segregating suppressors, we surveyed 90 strains from the *Drosophila* Genome Reference Panel. Rather than loss of *Rsp*, we found that over half of the strains (52/90) harbor suppressors located on the X chromosome or autosomes, but not the Y chromosome. The widespread presence of strong suppressors limited the resolution of our genome-wide association mapping; however, recombination analysis identified a strong X-linked suppressor to a ∼300 kb interval on the chromosome. Together, our findings suggest that pervasive, multilocus suppression constrains the spread of *SD* in natural populations.

## Introduction

Not all genetic factors benefit their hosts. Selfish genetic elements gain a transmission advantage to the next generation, often at the host’s expense. Meiotic drivers are a classic example — they disrupt Mendelian inheritance to increase their frequency among viable gametes [1]. Despite this selfish advantage, most meiotic drivers are typically observed at low frequencies in natural populations [2, 3]. For instance, *Segregation Distorter* (*SD*), one of the most well-studied meiotic drivers, is present in most populations of *Drosophila melanogaster* worldwide, but segregates at frequencies <10% in each population [4, 5]. The population dynamics of meiotic drivers, such as *SD,* can be influenced by their fitness costs, the frequency of sensitive target chromosomes, and the impact of other host genetic factors and environmental conditions [6–11].

*SD* biases its transmission by eliminating sperm carrying target sequences during spermatogenesis. The *SD* system is autosomal and includes two major components: the driver on chromosome 2L, a partial tandem duplication of *Ran GTPase Activating Protein* (*RanGAP,* referred to as *Sd-RanGAP*); [12] and its target, *Responder* (*Rsp*), in the pericentromeric heterochromatin on chromosome 2R. *Rsp* corresponds to a large block of tandem ∼120-bp satellite DNA repeats whose copy number positively correlates with sensitivity to *SD*. Only chromosomes with more than 100 repeats are susceptible to distortion [8, 13]. Besides *Sd-RanGAP* and *Rsp*, many other genetic factors can modify the drive strength of *SD*. *SD* chromosomes recruit enhancers to strengthen drive. We know of at least three enhancers of drive, including *E(SD)*, *M(SD),* and *St(SD)*, that exist on specific *SD* chromosomes. Loss of any of these enhancers reduces drive strength [14–17]. To maintain the linkage between *Sd-RanGAP*, insensitive *Rsp,* and enhancers of drive, *SD* chromosomes frequently recruit chromosomal inversions to suppress recombination [18]. While these inversions help preserve coadapted complexes, reduced recombination can lead to the accumulation of deleterious mutations [19–21]. The costs associated with *SD* through drive itself [6, 22] create selection pressure for the evolution of suppressors. Suppressors of *SD*, found on both the X chromosome and autosomes, prevent *SD* from killing sperm [10, 23]. Although these *SD*-modifying factors were identified decades ago, their molecular identities and natural dynamics remain unknown [11, 23–25].

To address this knowledge gap, we investigated the spectrum of *Rsp* alleles and the frequency of factors modifying drive strength using sequenced inbred lines from North Carolina, USA (Drosophila Genetic Reference Panel, DGRP;[26]). Our findings reveal a rarity of insensitive *Rsp* alleles and a high frequency of suppressors in this population, suggesting that pervasive suppression, instead of the absence of sensitive *Rsp*, halts the spread of *SD* chromosomes in the US. Leveraging genome-wide association studies (GWAS) and recombination mapping, we identified several genetic modifiers and discovered a strong X-linked suppressor. These results provide novel insights into the complex dynamics of *SD* chromosomes and their suppressors.

## Results

### Low frequency of insensitive *Rsp* in the DGRP

We first investigated whether the low frequency of *SD* in North America could be explained by the loss of its target, *Rsp* [5, 10, 27]. Given the inherent challenges in quantifying repetitive sequences [28–30], we employed three orthogonal approaches to estimate *Rsp* copy number in 86-90 DGRP lines: one computational approach using DGRP Illumina DNA sequence data, and two molecular approaches based on slot blots and quantitative PCR (qPCR; Fig. 1A; Table S1). Each approach yields the relative abundance of *Rsp* repeats. We estimate copy number by comparing it to the reference strain (*Iso-1*; [31]). Results from all three quantification methods are highly correlated (Spearman’s rho ≥ 0.63; Fig. 1B), indicating that each approach robustly captures *Rsp* copy number variation.

**Fig 1.**
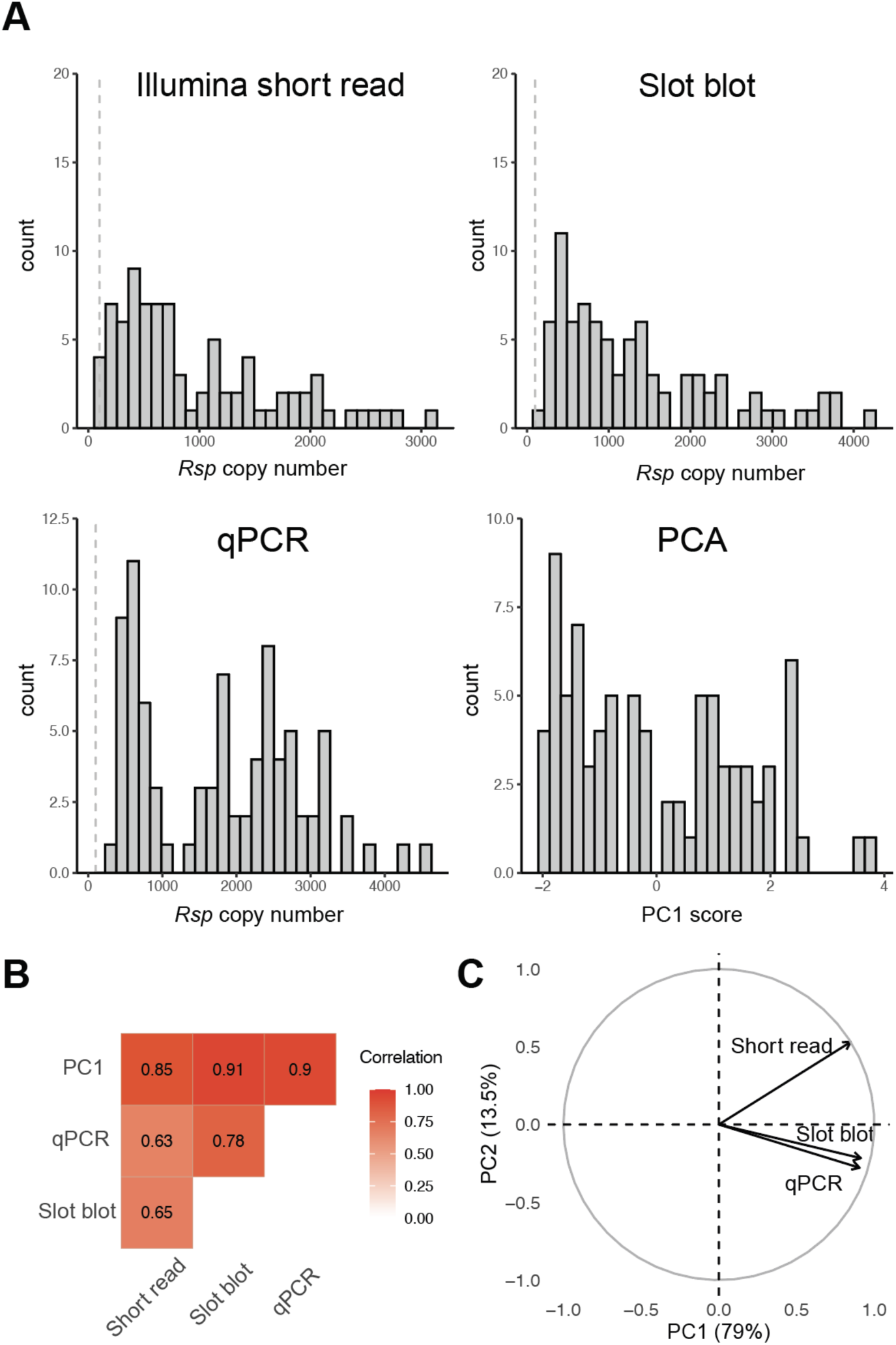
Distribution of *Rsp* copy number in the DGRP. **(A)** Histograms showing the distribution of *Rsp* copy number as measured by Illumina short-read sequencing, slot blot, and qPCR, alongside the distribution of scores for the first principal component (PC1). The dashed line represents 100 copies of *Rsp*, and chromosomes carrying less than 100 copies of *Rsp* are likely insensitive to *SD* drive. **(B)** Pairwise correlations of *Rsp* copy number between the three estimation methods and PC1. Each method is strongly correlated with the other, indicating they can robustly estimate the *Rsp* copy number. The number shows Spearman’s correlation coefficients. **(C)** A PCA variable loading plot showing the contribution of each measurement technique to the first two principal components. The percentage of total variance explained by PC1 (79%) and PC2 (13.5%) is indicated on the axes.

Nevertheless, each method carries its own biases. Computational approaches may be affected by library construction artifacts and reference alignment bias. Slot blots are more sensitive to variants that closely match the probe sequences, whereas qPCR can be influenced by differences in amplification efficiency and primer specificity. To mitigate method-specific biases, we integrated the three estimates using principal component analysis (PCA). The first principal component (PC1), which explained 79% of the total variance, was used as a composite proxy for *Rsp* copy number variation (Fig. 1C; Table S1). In subsequent analyses, we also reported results using individual estimation methods to assess the accuracy of each estimation and the robustness of our conclusions.

Most DGRP lines appear to carry *Rsp* alleles sensitive to *SD*, with a mean of 946, 1355, and 1782 copies using the computational, slot blot, and qPCR methods, respectively (Fig. 1A). These values are comparable to the *Iso-1* reference allele, which carries ∼1100 *Rsp* repeats from the US population. The discrepancy among methods could in part be due to the underrepresentation of pericentromeric repeats in Illumina data, or changes in *Rsp* copy number over the ∼15 years since those sequences were generated. Two lines, RAL28 and RAL73, had fewer than 100 *Rsp* repeats in the computational methods, but not in other methods, making them likely to be insensitive or intermediate-sensitive to *SD* [8, 13, 15]. Our results indicate that, despite the cost of sensitive *Rsp* when *SD* is present in the population, some form of selection may maintain high *Rsp* copy numbers.

### High frequency of suppressors in the DGRP

Despite the prevalence of *Rsp* alleles with high copy numbers, which should render them sensitive to *SD*, the frequency of *SD* remains low. This suggests that many strains may evade distortion by carrying suppressor alleles that mitigate the effects of *SD*. To survey natural variation in resistance to an *SD* chromosome in the DGRP, we used an *SD* chromosome from Madison, WI (*SDMad*; [32]), containing the dominant marker *Curly* (*Cy*), that exhibits strong drive [33]. Using reciprocal crosses between DGRP strains and *Cy SDMad* (Fig. 2A), we assessed the sensitivity of each DGRP strain to drive. One cross generated F1 males with an X chromosome from the DGRP stock (DGRP-X), while the other generated F1 males with the X chromosome without suppressors from the *Cy SDMad* stock, which lacks suppressors (No X Suppressors). We then crossed individual F1 males to 2 females and generated 5 to 10 single biological replicates for each genotype to estimate their drive strength. We quantified drive strength by calculating proportion of offspring with *SD* (*k* value). The expected value of *k* in the absence of drive is 0.5. Without suppressors, *Cy SDMad* induces strong drive over wild-type alleles and produces offspring only carrying itself (*k* value = 1.0). These crosses revealed significant variation in sensitivity to *SD* across the DGRP (Fig. 2B; Table S2), but minimally within strains. Therefore, in the following analyses, we merged progeny counts from all replicates and calculated a single representative *k* for each genotype.

**Fig 2.**
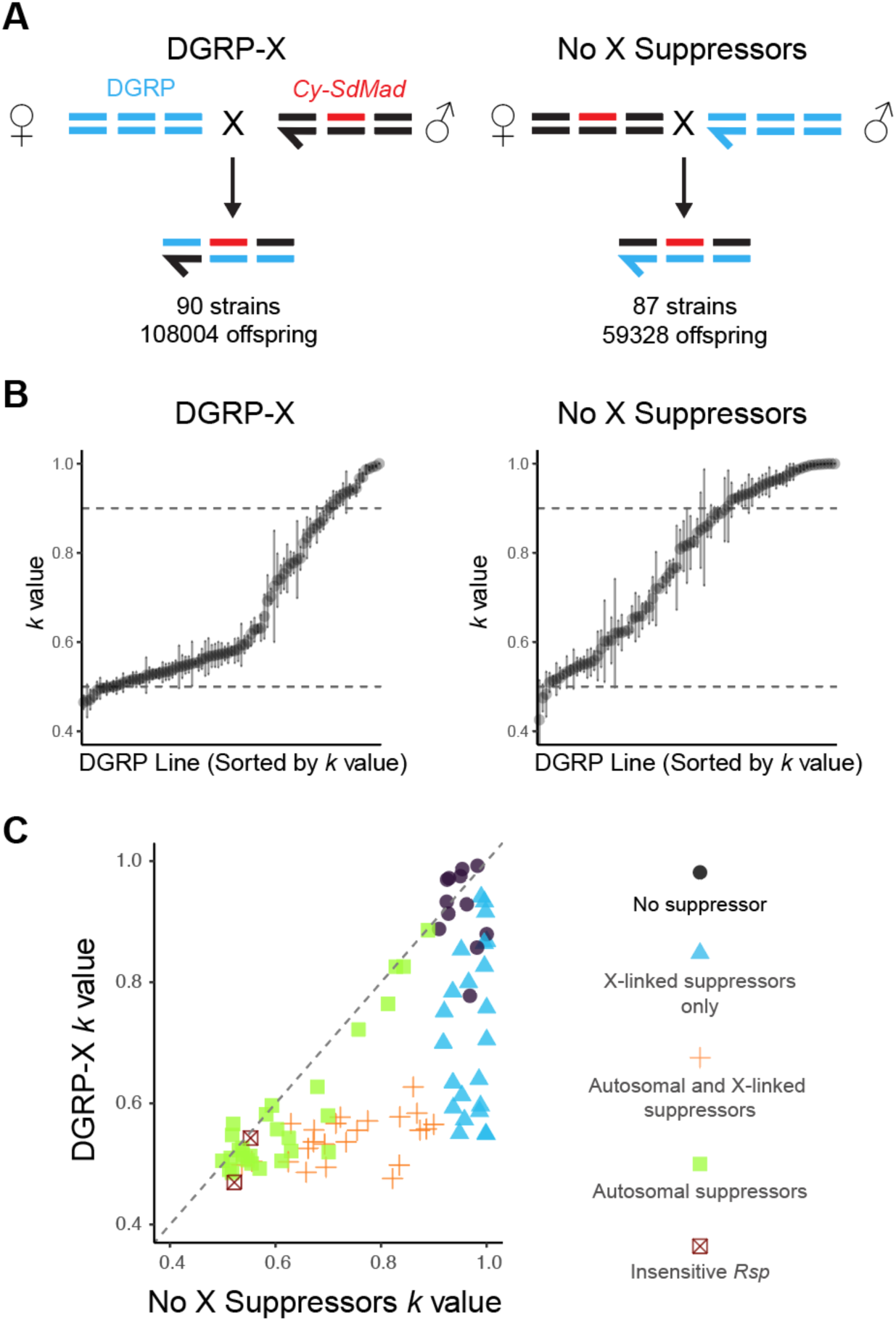
High frequency of suppressors on autosomes and the X chromosome across DGRP strains. **(A)** To assess the strength of segregation distortion (measured as *k* value: proportion of offspring with *SD*), we crossed each DGRP strain with a strain carrying an *SD* chromosome with a dominant marker, *Cy (Cy-SdMad)*. In the DGRP-X panel (left), DGRP females were crossed to males carrying the *SD*, followed by test crosses of F1 males to control females. In the “No X Suppressors” panel (right), we generated F1 males without X-linked suppressors from reciprocal crosses. We indicate the number of DGRP strains and offspring surveyed for each cross. **(B)** Distributions of *k* values across DGRP lines for each panel, with lines sorted by mean *k* value. Each point represents the mean *k* value ± standard error. Dashed lines mark the thresholds for the drive without suppressors (*k* > 0.9) and no drive (*k* = 0.5). **(C)** Scatterplot comparing *k* values from the DGRP-X and No X Suppressors and panels for each DGRP line. We inferred the presence of autosomal suppressors if the k value in the "No X Suppressors" panel was less than 0.9 (excluding two strains, RAL28 and RAL73, which are likely Rsp-insensitive with <100 Rsp repeats). We inferred the presence of X-linked suppressors if the k value from the DGRP-X panel was significantly less than that from the "No X Suppressors" panel (Student’s t-test, P < 0.05). Colors and shapes indicate the inferred presence and position of suppressors in each strain: no suppressors (black circles), X-linked suppressors only (blue triangles), both X-linked and autosomal suppressors (orange crosses), autosomal suppressors (green squares, which might mask the presence of X-linked suppressors), and *Rsp*-insensitive lines (red X). The diagonal dashed line represents equal distortion in both crosses.

As expected [7], *Cy SDMad* does not strongly drive (*k*<0.56; Fig 2C) in crosses involving the two strains (RAL28 and RAL73) with putatively insensitive *Rsp* alleles. For the rest of the 88 strains carrying sensitive *Rsp*, the “DGRP-X” crosses allowed us to estimate the overall frequency of suppressors located on any chromosome. Using a threshold of *k* < 0.9 to indicate the presence of suppressors (see Materials and Methods for justification), we found that 74 DGRP strains (84%) carry factors that suppress drive. In contrast, the “No X suppressors” cross design excludes the contribution of DGRP X chromosomes, thereby revealing that 51 DGRP strains (58%) carry dominant autosomal suppressors or less sensitive *Rsp* (Fig. 2C).

To confirm our inference that there are dominant suppressors on autosomes, we randomly isolated nine second chromosomes and seven third chromosomes from the DGRP strains with evidence for autosomal dominant suppressors (Fig. S1A). Among these strains, six second chromosomes from RAL38, RAL59, RAL45, RAL324, RAL707, and RAL375 and three third chromosomes from RAL59, RAL357, and RAL324 have ability to suppress *SD* (*k* < 0.9; Fig. S1B). Based on these data, we estimated 39% of second chromosomes carry dominant suppressors or intermediate sensitive *Rsp* and 25% of third chromosomes carry dominant suppressors in the population.

To infer the presence of sex-linked suppressors, we directly compared the results from reciprocal crosses. We observed that 46 strains showed significantly higher drive in F1 males from “No X suppressors” than from “DGRP-X”, indicating the presence of X-linked suppressors (single-tail Student’s t test, P<0.05; Fig. 2C). Among them, 39 X-linked suppressors decreased *k* by more than 0.1. This is a conservative estimate, as strong autosomal suppressors could mask the effects of X-linked ones. In contrast, for all tested DGRP strains, drive strength was equal or stronger when F1 males inherited a DGRP Y chromosome (“No X suppressor”), providing no evidence for Y-linked suppressors (Fig. 2C). In conclusion, our results suggest that X-linked suppressors are present at relatively high frequencies in the DGRP population (>52%), which may contribute to the low frequency of *SD* chromosomes observed in the US [10].

### *SD* drive strength is influenced by the interplay of *Rsp* copy number and suppressors

To map the modifiers of drive strength, we performed genome-wide association studies (GWAS) using DGRP sequence information. We first mapped Illumina reads to the latest R6 genome assembly, which allowed us to call single-nucleotide polymorphisms (SNPs) more accurately than analyses based on the earlier *D. melanogaster* assembly (R5), which lacked coverage of many heterochromatic regions. After filtering low-frequency (<10%) SNPs and missing (>10%) SNP data among strains, we retained 1,283,850 SNPs for GWAS analyses.

*Rsp* copy number correlates positively with sensitivity to drive [8, 34]. We examined the relationship between *Rsp* copy number and *k* values and found a strong positive correlation across all estimation methods (R = 0.75 and 0.61 for PC1; Fig. 3A and Fig. S2). These results not only support our estimates of *Rsp* copy number variation but also indicate that, even in the presence of suppressors, *Rsp* copy number remains a key determinant of *SD* drive strength.

**Fig 3.**
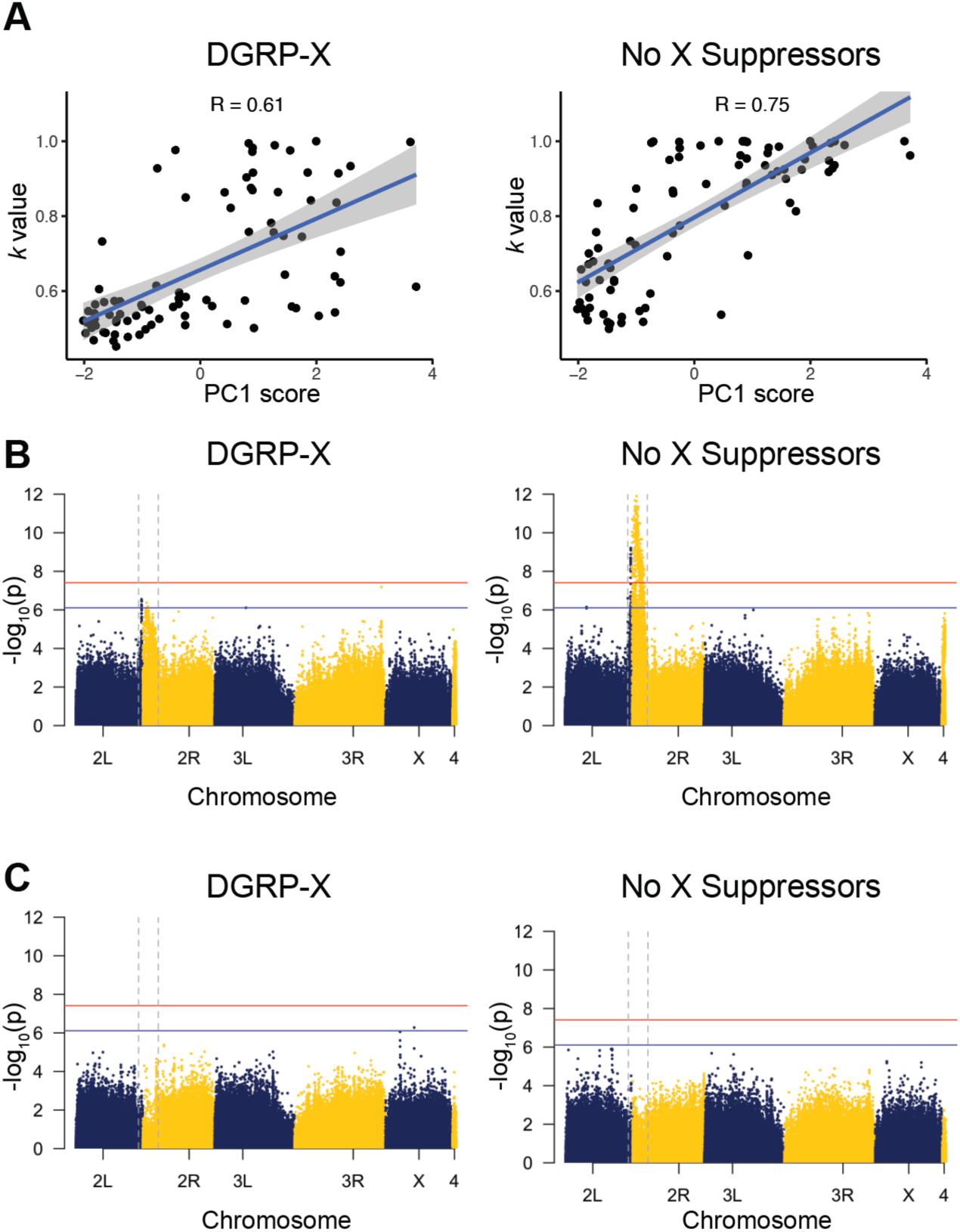
*Rsp* copy number is associated with drive strength and explains GWAS signals near the centromere of chromosome 2. **(A)** Scatter plots showing the correlation between drive strength (*k*) and the first principal component (PC1) summarizing *Rsp* copy number estimates across DGRP strains. The left panel shows the result from F1 males carrying the X chromosome from DGRP (DGRP-X), and the right panel shows the result from the reciprocal cross (no X suppressors). The high correlation coefficient (R) in each panel suggests that second chromosomes with higher *Rsp* copy are more sensitive to *SD*, even in the presence of suppressors. **(B)** Manhattan plots of GWAS results for *k* in the DGRP-X panel (left) and in the “no X suppressors” (right). Each dot represents a SNP, and chromosomes are shown. The red and blue horizontal lines denote genome-wide significant and suggestive thresholds after Bonferroni’s correction. We observed a strong signal near the centromere on chromosome 2 (between vertical dashed lines). **(C)** Manhattan plots as in (B), but with PC1 of *Rsp* copy number included as a covariate in the GWAS model. The signal near the centromere on chromosome 2 is substantially reduced, suggesting that *Rsp* copy number accounts for the association. Quantile-quantile plots are in Fig. S4.

Our initial GWAS analyses, without incorporating the effect of *Rsp* copy number, revealed a broad peak spanning the centromere of chromosome 2 (Fig. 3B; Fig. S3A). This signal may reflect SNPs linked to sensitive *Rsp* loci or enhancers, as two known enhancers— *E(SD)* and *M(SD)*— are located in the pericentric heterochromatin of chromosome 2 [17]. To account for the impact of *Rsp*, we included our estimated number of *Rsp* repeats for each strain as a covariate in our analyses. This adjustment significantly attenuated the association signals near the centromere of chromosome 2 (Fig. 3C and Fig. S3-4). Notably, when PC1 (composed of three *Rsp* estimates) was included as a covariate, the GWAS signal in this region was no longer detectable, indicating that the association might be primarily driven by *Rsp* copy number (Fig. 3C).

Incorporating *Rsp* copy number estimates into the GWAS also allowed us to detect a significant SNP on chromosome 2L. However, we did not identify any clear candidate loci on the other chromosomes (Fig. 3C), despite evidence from genetic crosses suggesting the presence of X-linked suppressors at high frequency (>52%). One possible explanation is that the presence of multiple strong suppressors within our sample could be masking the effects of individual suppressors, making them difficult to detect. Additionally, our relatively small sample size compared to other DGRP-based GWAS likely reduces our statistical power to detect associations on the X and autosomes.

### Recombination mapping of a strong X-linked suppressor

To overcome the challenges with identifying suppressors using GWAS analyses, we focused on one strong *Su(SD)X* chromosome from RAL256 and asked how many major genetic loci contribute to the suppressing effect. We generated 239 recombinant X chromosomes between *Su(SD)X* and a wildtype X chromosome with a *y* marker from *Iso-1*, and measured the degree of suppression (Fig. 4A; Table S3). The bimodal distribution of *k* values among recombinants, with peaks at *k* = 0.55 and 1.0, suggests the presence or absence of a major suppressor. By using a threshold of *k* < 0.9 to define suppression, we partitioned the recombinant population into two groups. Approximately 54% of recombinant X chromosomes are capable of suppressing SDMad (*k* < 0.9), indicating that a single major locus, hereafter referred to as *Su(SD)XM*, is the primary contributor to this suppression effect (Fig. 4B). We further inferred that the *Su(SD)XM* randomly segregates with the visible marker *y* (133/239; 56%, Fisher exact test’s P = 0.09), indicating that *Su(SD)XM* is located away from the X chromosome telomere, where *y* resides(Fig. S5).

**Fig 4.**
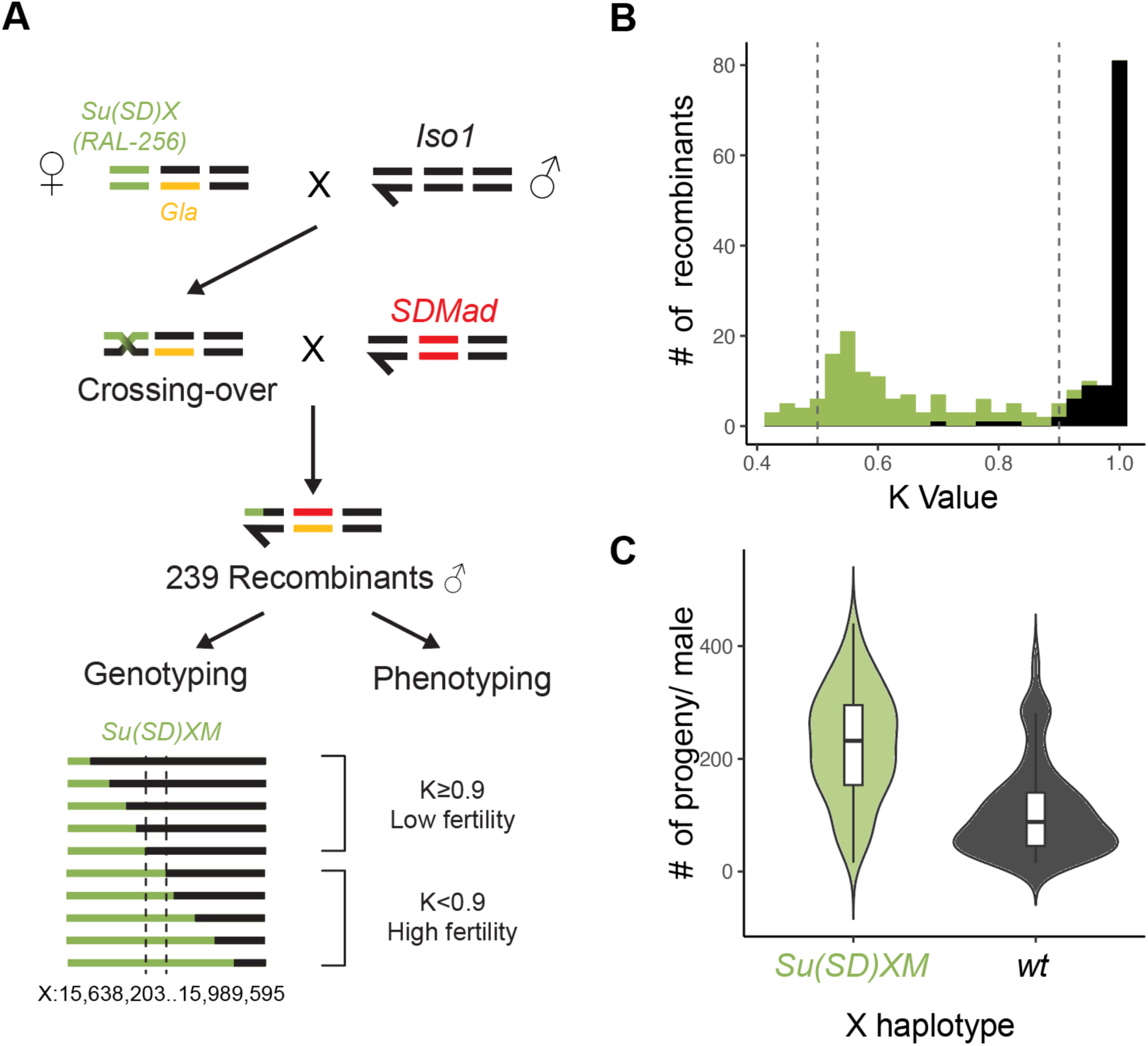
One major locus contributes to the suppressing effect of *Su(SD)X* by rescuing sperm with sensitive *Rsp* alleles. **(A)** We generated 239 recombinants between the *Su(SD)X* from RAL-256 and *y* chromosomes. We measured *k* values of these recombinants by counting their offspring. We also genotyped these recombinants to map suppressors. The recombining chromosomes containing 350 kb from the *Su(SD)X* chromosome (marked in green) can suppress the *SDMad* chromosome **(B)** and produce more offspring **(C)**.

To map the candidate region for *Su(SD)XM*, we designed 15 primer pairs based on the indels between *Su(SD)X* and the *y* chromosome to genotype recombinant X chromosomes (Table S4). Based on our mapping results, we narrowed down *Su(SD)XM* to the region close to X:15,638,203..15,989,595, in R6 coordinates, which contains 88 genes (Fig. 4A; Table S3). This region is located close to one identified X-linked suppressor from *lt stw*^3^ stock (between *yellow* and *carnation,* X: 356,509 (y) - 19,569,401)[35] and another one in a lab strain (between 13C7 and 13E4; X: 15,506,633-15,670,816)[36]. However, we did find four recombinants that might not harbor *Su(SD)XM* but have *k* values of 0.697, 0.786, 0.789, and 0.897. Vice versa, three recombinants that might harbor a suppressor have *k* values of 0.901, 0.919, and 0.920, respectively (Fig. 4B). This inconsistency may correspond to stochastic variation in drive strength [14], *de novo* mutations affecting drive strength of the *SD* chromosome in our recombinants [33], or imperfect environmental control [11].

Additionally, *SD-*bearing males with *Su(SD)XM* produced about two-fold more offspring than the males without *Su(SD)XM* (225 vs. 104 offspring per male; Student’s t-test’s P <0.001 Fig. 4C; Table S3). This result suggests that *Su(SD)XM* can directly rescue sperm from being killed by *SD*, making the suppressor advantageous when *SD* is segregating in a population. If other suppressors similarly improve male fertility in the presence of *SD*, the high frequency of suppressors in the DGRP may be a direct outcome of counteracting *SD*.

To identify SNPs associated with the *Su(SD)XM* suppressor, we performed a haplotype-based analysis on the DGRP population. Compared to *Iso-1,* the reference strain without *Su(SD)XM*, RAL256 carries 10,249 SNPs nearby or within the identified *Su(SD)XM* region (X:15,500,000..16,000,000). Among them, 4,983 are absent from all suppressor-free strains, which are likely tightly linked with *Su(SD)XM.* However, none of these 4,983 SNPs were present in more than half of the 39 strains inferred to have X-linked strong suppressors. This result indicates, in addition to *Su(SD)XM*, other distinct X-linked suppressors contribute to the resistance of *SD* in the DGRP population.

## Discussion

The *Segregation Distorter* (*SD*) chromosome promotes its transmission by killing sperm bearing sensitive alleles of a target that *SD* chromosomes lack. Given this powerful advantage, a long-standing puzzle is why *SD* chromosomes are so rare in the wild [37, 38]. In this study, we surveyed and mapped modifiers of *SD* chromosomes in a sequenced inbred population (DGRP; [26]). We found that the suppressors of *SD* are segregating at a high frequency, with only two of 90 strains carrying insensitive *Rsp* alleles. This high frequency of suppressors, together with the known deleterious effects of *SD* chromosomes [6], likely explains the low frequency of *SD* chromosomes in the US population [5, 36, 39, 40].

Our observation of very few insensitive *Rsp* alleles in non-SD chromosomes suggests that mutating the *SD* chromosome’s target, *Rsp*, may not be an optimized long-term solution, as *Rsp* deletions may result in reduced fitness [13]. Although such *Rsp* deletions can be favored when *SD* chromosomes segregate in a population lacking suppressors [13], their advantages diminish once suppressors emerge. After suppressors become common, *Rsp* repeats may rapidly recover their copy number via unequal crossing-over and selection for high *Rsp* number. Such evolutionary turnover, triggered by *SD* or other meiotic drivers, could further promote the rapid evolution of repeats across species.

While our initial GWAS screen for suppressors failed to detect any on the X chromosome, a more focused recombination mapping technique revealed that a potent suppressor was indeed present. This suggests that the phenotype is controlled by multiple genetic factors and their epistatic interactions. The effects of strong suppressors can be masked by other suppressors or by the number of *Rsp* repeats, as we did not detect evidence of negative distortion (where *k <* 0.5) caused by multiple strong suppressors in the population (Fig. 2). This non-additive effect makes modifiers invisible to certain analytical tools and highlights the intricate, layered nature of this genetic conflict.

Intriguingly, we identified a hotspot for these suppressors on the X chromosome, where they are found at a higher frequency (>52%) compared to other chromosomes (<40%). This pattern can be partly explained by detection bias: while our screen only detects dominant autosomal suppressors, any X-linked suppressor, whether dominant or recessive, is revealed in males due to their hemizygosity. In addition, for the same reason, X-linked suppressors are fully exposed to selection and may increase in frequency more rapidly, regardless of dominance [41]. Together, these factors suggest that the observed enrichment of X-linked suppressors may be a result of a combination of detection and selection biases.

This raises another unresolved question: why does this population maintain such a large and diverse arsenal of suppressors against *SD* chromosomes with the same driver? This pattern may reflect the fact that suppressors vary in their effectiveness depending on the specific *SD* chromosomes present [10, 23, 24, 36, 42]. Given that multiple *SD* chromosomes carrying different inversions are segregating in the US populations [5, 36, 39, 40], it is likely that different *SD* chromosomes require distinct suppressors, maintaining multiple suppressors in the population.

Additionally, modifiers may be relatively easy to evolve and may even arise from pre-existing standing genetic variation. This idea is supported by evidence that disruption of many genes in different pathways can alter *SD* drive strength [43–45]. Furthermore, suppressors exist at high frequencies in some populations that lack *SD* chromosomes, such as in Texas [10], though not in Japan [21]. Similarly, studies on sex-ratio meiotic drive in other systems suggest that suppressive mechanisms can emerge from both new mutations, such as the evolution of novel siRNA genes [46–48] and standing variation (e.g. [49]). Identifying the specific origins of *SD* suppressors remains a key question for future research.

Genetic conflicts induced by meiotic drive systems often exploit fundamental cellular processes [50, 51]. For example, the primary driver of *SD* is a Ran GTPase-activating protein, which acts as a nuclear transport factor in the Ran signaling pathway [12]. The range of potential mutations available for the evolution of drive modifiers is therefore likely to be broad [52]. These evolutionary arms races, where both drivers and suppressors can rise in frequency and even become fixed in populations despite their fitness costs, can accelerate the evolution of genes involved in spermatogenesis and chromosome segregation across diverse species [53–57].

## Materials and Methods

### Assaying drive strength of *Segregation distorter (SD)*

We measured drive strength of a marked *SD* chromosome (*Cy SDMad* [33]) as the proportion of *SD* offspring (*k*) in crosses with different genetic backgrounds. For surveying drive strength in DGRP strains, we used two independent cross schemes (Fig. 2A). For recombination mapping of *Su(SD)X,* we used the crossing scheme in Fig. 4A to generate and test drive strength in males with recombinant X chromosomes with the marked *Cy SD* chromosome. We crossed one 0 to 5-day-old male to two *Iso-1* females and transferred every 4 days for either 20 days (DGRP crosses) or 12 days (recombination mapping). We counted all offspring produced from 5–10 individual males for each strain, and only replicates with more than 40 offspring were reported and used in the following analyses. We measured drive strength (*k* values) by the number of offspring with *Cy* or SdMad over the total number of offspring [5].

### Estimating *Rsp* copy number in DGRP using Illumina short reads

We mapped Illumina reads from DGRP [26] and *Iso-1* to the reference from Chang [58, 59] using Bowtie2 with the --fast preset (v2.3.5.1; [60]). We used a Python script (htseq_bam_count_proportional.py; https://github.com/LarracuenteLab/Dmelanogaster_satDNA_regulation) to count reads that mapped to *Rsp* and estimate relative counts as reads per million (RPM) values [61]. We estimated the *Rsp* copy number in each DGRP strain based on the number of reads mapped to the *Rsp* locus and 1,100 copies of *Rsp* in *Iso-1* [62]. We assume that the number of reads mapped to the *Rsp* locus is linearly correlated with *Rsp* copy number.

### Estimating *Rsp* copy number in DGRP using slot blots

The *Responder* biotin probe was generated using previous methods [31]. Briefly, RNA was made in a 40ul T7 transcription reaction with 400ng of PCR template, amplified from a set of cloned Responder repeats, and biotin label mix (Roche 11685597910), and the RNA was isolated using Monarch RNA cleanup kit (New England Biolabs T2050S).

We mixed 600ng of each genomic DNA with 140ul denaturing buffer (0.5N NaOH, 500mM NaCl), incubated at room temperature for 10 minutes, and then mixed with 150ul ice-cold loading buffer (1x SSC, 0.125N NaOH). One hundred microliters of each denatured DNA (200ng) was loaded onto a wetted Biodyne B 0.45um Nylon membrane [Thermo Scientific 77016] in triplicate using Bio-Dot SF slot blotter [Bio-Rad #1706542], and each slot was rinsed with 150ul loading buffer and 200ul 400mM Tris-Cl (pH 7.5). After rinsing the membrane with 2x SSC (300mM NaCl, 30mM sodium citrate), it was cross-linked to the membrane with UV light. After the membrane was incubated with 12ml ULTRAhyb Ultrasensitive hybridization buffer [Invitrogen AM8670] for 1 hour at 42 °C, 1.5 μL 100 μM sec5-Rp49 oligos and 3 μL 100 μM sec5-IRD800 oligo (IRD880-AACACCCTTGCACGTCGTGGA) were added and hybridized at 42°C overnight. After washing the membrane 3 times with 2xSSC/ 0.1% SDS at 42°C for 15 minutes each wash, the membrane was imaged with the 800nm channel and quantified on a LI-COR Odyssey CLx Imager.

The membrane was incubated with 12ml hybridization buffer at 42°C for 1 hour, 500ng Responder biotin probe added, and incubated at 42°C overnight. After washing the membrane 4 times with 2xSSC/0.1% SDS at 42 °C for 15 minutes each wash, Chemiluminescent Nucleic Acid Detection Module [Thermo Scientific 89880]; the membrane was incubated in 16ml Blocking buffer with 55ul streptavidin-HRP conjugate for 15 minutes at room temperature. After three washes with 15 mL of 1x Block Wash buffer at room temperature for 5 minutes. The membrane was incubated with the enhancer/peroxide solution for 5 minutes at room temperature, and imaged and quantified on the Bio-Rad Chemi-Doc XRS+. Each membrane contained a triplicate of Iso1 genomic DNA and the Responder repeat (chemiluminescence) signal of each slot was normalized to its Rp49 probe (IRD800) signal and the ratio for each line was normalized to the *Iso1* ratio and multiplied by 1100 (estimated *Rsp* copy number). *Rsp* number estimated from slot blotting is in Table S1.

### Estimating *Rsp* copy number in DGRP using qPCR

Serial dilutions of *Iso1* genomic DNA (190ng, 19ng, 1.9ng, 0.19ng, and 0.019ng) as a standard and 1.9ng genomic DNA from each line were assayed in triplicate in 15ul qPCR reaction using 2xSSO-Advanced master mix (Bio-Rad 172-5270) and 500nM final concentration of each primer (*Rsp* forward: GGAAAATCACCCATTTTGATCGC, *Rsp* reverse: CCGAATTCAAGTACCAGAC, tRNA forward: CTAGCTCAGTCGGTAGAGCATGA, tRNA reverse: CCAACGTGGGGCTCGAAC). The reactions were amplified on a Bio-Rad CFX96 machine (95 °C 30 seconds, 40 cycles of 95 °C 10 seconds, 60 °C 10 seconds). The *Rsp* and tRNA (Lys-CTT) copy numbers for each line were quantified by comparing their threshold cycle to the *Iso1* standard curves created using the serial dilutions, using 1100 as the *Iso1 Rsp* copy number. *Rsp* number estimated from qPCR estimation is in Table S1.

### Principal component analysis of *Rsp* copy number

We performed PCA on an 86-sample subset where estimates were available for all three *Rsp* copy number estimations. We ran PCA in R using the prcomp() function on centered and scaled copy number data. When necessary, the principal component scores were negated to obtain a positive correlation with *k*.

### GWAS analyses using updated SNP calls

We applied the GATK4 best practice pipeline to map and call SNPs in DGRP [63] using the R6 genome assembly [62]. In short, we mapped Illumina reads from DGRP and iso-1 to the Flybase R6 reference using bwa mem (v0.7.15; [64]) and marked the duplicates using GATK MarkDuplicates with default settings (v4.1.2; http://broadinstitute.github.io/picard/). We integrated known indel data from the DGRP [26] and DPGP1 [65] to optimize our SNP calling in GATK. Using the resulting SNPs, we conducted GWAS analyses using plink v1.90b7.2 [66] with parameters “--linear interaction --no-sex --allow-extra-chr --maf 0.1 --geno 0.1.” We also integrated *Rsp* copy number as covariates in our GWAS analyses. We calculated the genome-wide level of significance using a Bonferroni correction on the number of SNPs included in the analysis. A suggestive threshold was calculated using the same method with an alpha of one. We used the R package “qqman” [67] to make Manhattan and quantile-quantile plots.

### Recombination mapping of *Su(SD)X*

We genotyped 239 recombinants using PCR and electrophoresis on 2% agarose gels. The genotypes and phenotypes are listed in Table S3, and primer information is listed in Table S4.

## Acknowledgements

This work was supported by NIH R35 GM119515 and University of Rochester funds to AML. A.M.L. thanks Nathaniel and Helen Wisch for their support. C.-H.C. was supported by the Messersmith Fellowship from the University of Rochester and the Government Scholarship to Study Abroad from Taiwan. We thank Dr. David Houle and Dr. Jim Fry for their comments on this paper, and the Larracuente lab members and Dr. Daven Presgraves and Dr. Cara Brand for discussion. We also thank the University of Rochester CIRC for access to computing cluster resources.

## Supplementary Figures

**Fig S1.**
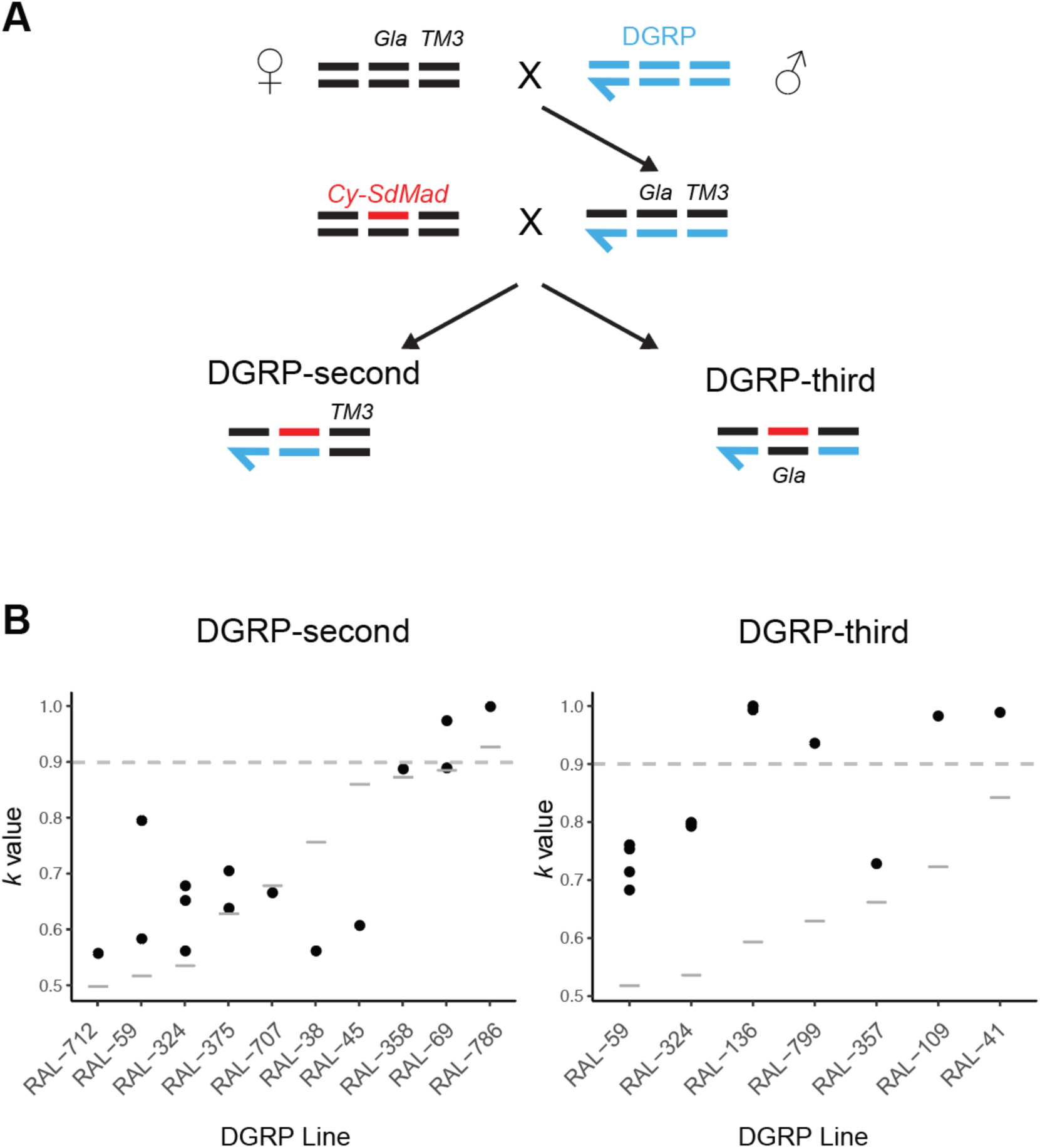
Surveys of dominant suppressors or intermediate sensitive *Rsp* on individual autosomes form DGRP. **(A)** Crossing scheme to introgress DGRP second or third chromosomes into a standard SD background. Males from each DGRP line were crossed to *Gla TM3* females, and F1 progeny carrying *Gla TM3* were crossed to *Cy-SdMad* tester flies to generate DGRP second or third chromosomes in a common *SD* background. **(B)** Variation in drive strength (*k* values) among DGRP lines for the second (left) and third (right) chromosomes. Black points indicate individual replicate measurements, and gray bars represent *k* from the “No X suppressors” crosses, indicating the combination of effects from all autosomes. Other than RAL-786, each DGRP strain tested here carries autosomes with dominant suppressors from previous surveys.

**Fig. S2.**
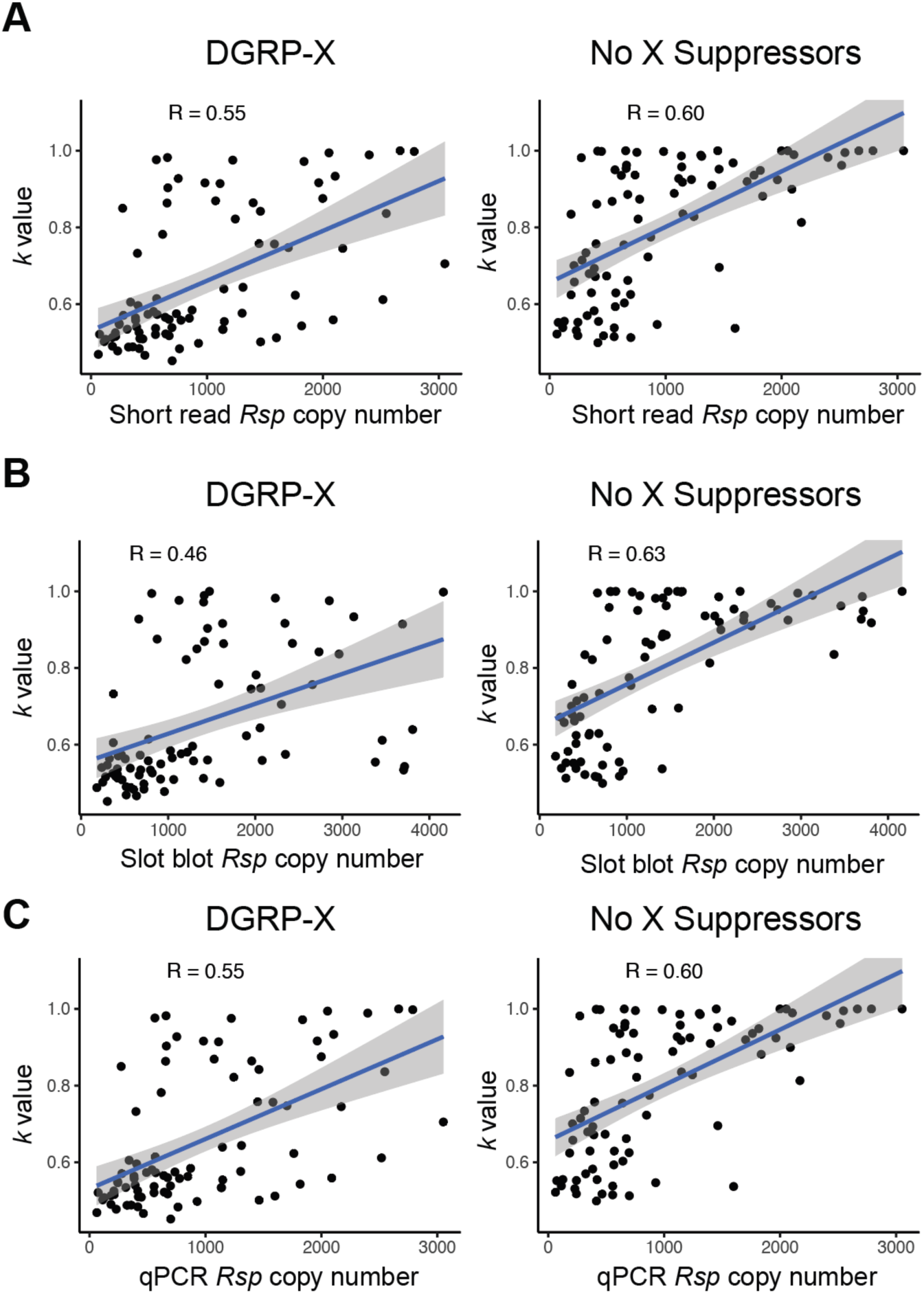
The positive correlation of *k* values and *Rsp* numbers using different methods across DGRP strains. There is a significant correlation between drive strength (*k* value) and Rsp copy number estimated from either Illumina short read sequence **(A)**, slot blot **(B)**, or qPCR **(C)**. The left panel shows the result from F1 males carrying the X chromosome from DGRP (DGRP-X), and the right panel shows the result from the reciprocal cross (no X suppressors). The high correlation coefficient (R) in each panel suggests that second chromosomes with higher *Rsp* copy are more sensitive to *SD*, even in the presence of suppressors.

**Fig. S3.**
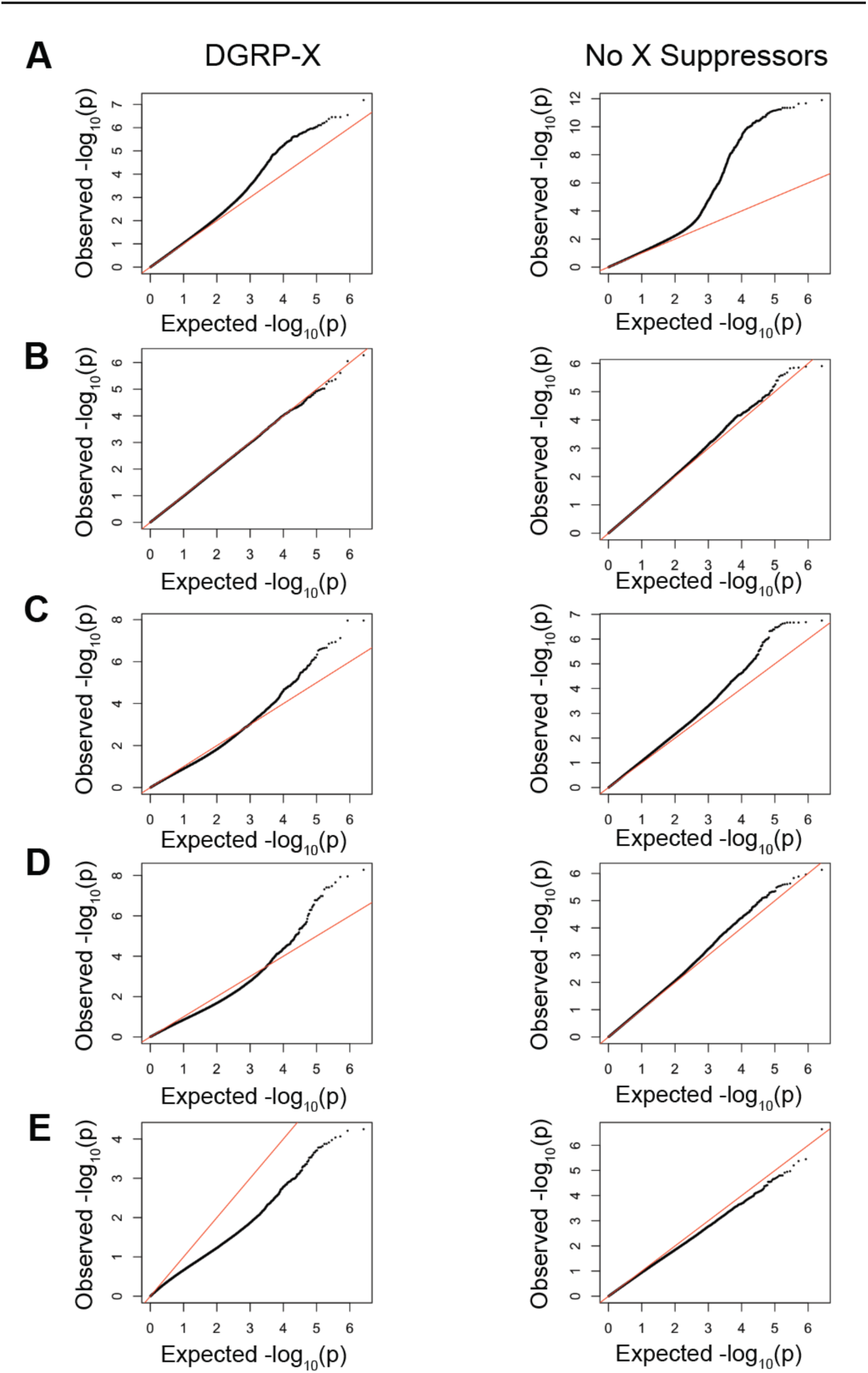
Quantile-quantile (Q-Q) plots from genome-wide association studies using DGRP strains under different conditions. We generated Q-Q plots to examine the distribution of observed p-values against expected p-values under the null hypothesis for GWAS results. Each row represents using covariates from different methods, including no covariate **(A),** PC1 **(B)**, Short reads **(C),** slot blot **(D),** and qPCR **(E).** The left panel presents the results for the "DGRP-X" crossing and the right panel presents the results from the "No X Suppressors" crossing. The black points represent the observed data, the red line indicates the expected distribution under the null hypothesis (y=x), and deviations from this line suggest potential population stratification or true association signals.

**Fig. S4.**
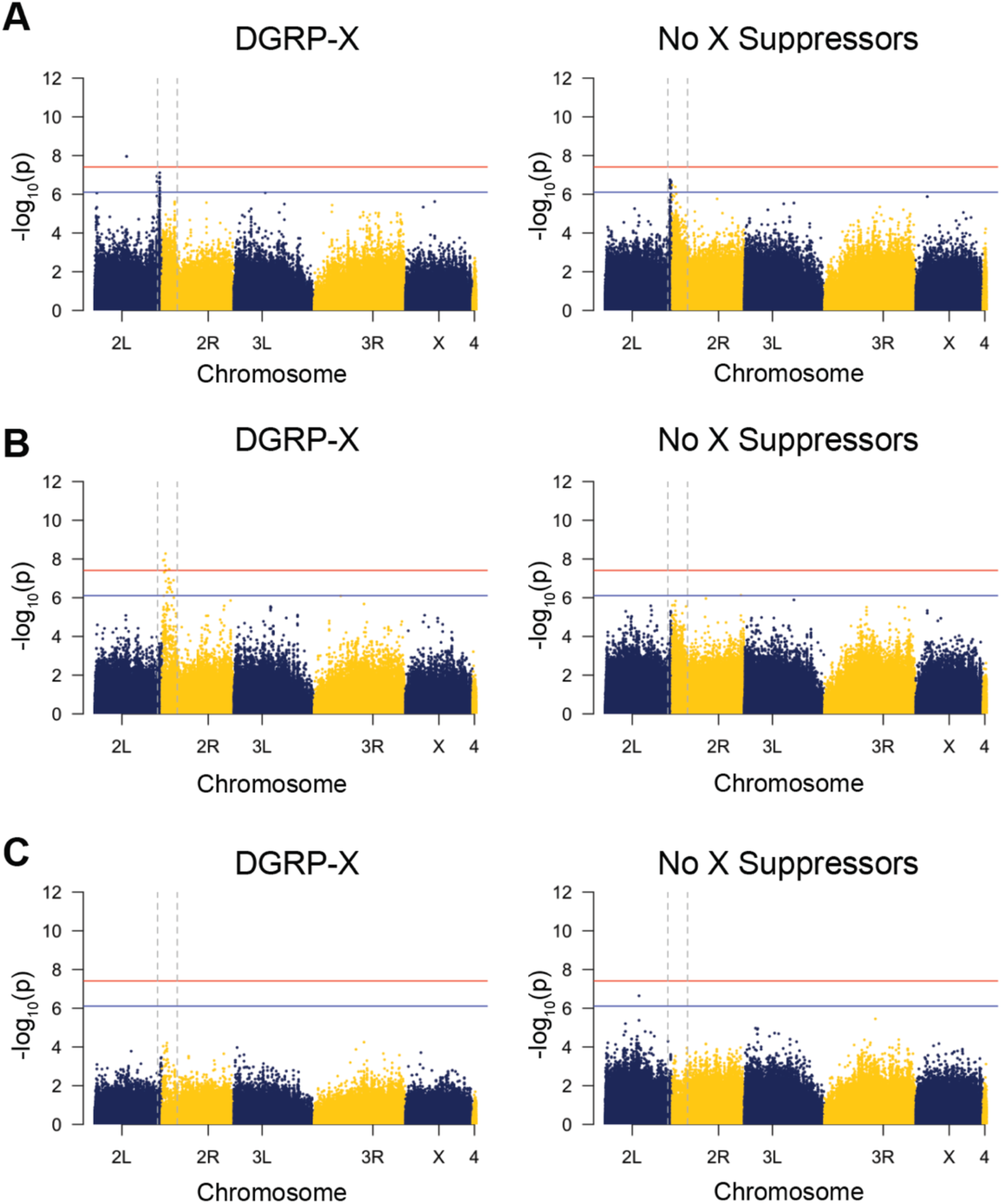
Manhattan plots of GWAS results for drive strength (*k*) using covariates from different *Rsp* estimation methods. We used *Rsp* copy number estimation from short read **(A)**, slot blot **(B)**, and qPCR **(C)**. Results using *k* from the DGRP-X panel (left) and from the “no X suppressors” (right) are shown, and each dot represents a SNP. The red and blue horizontal lines denote genome-wide significant and suggestive thresholds after Bonferroni’s correction.

**Fig S5.**
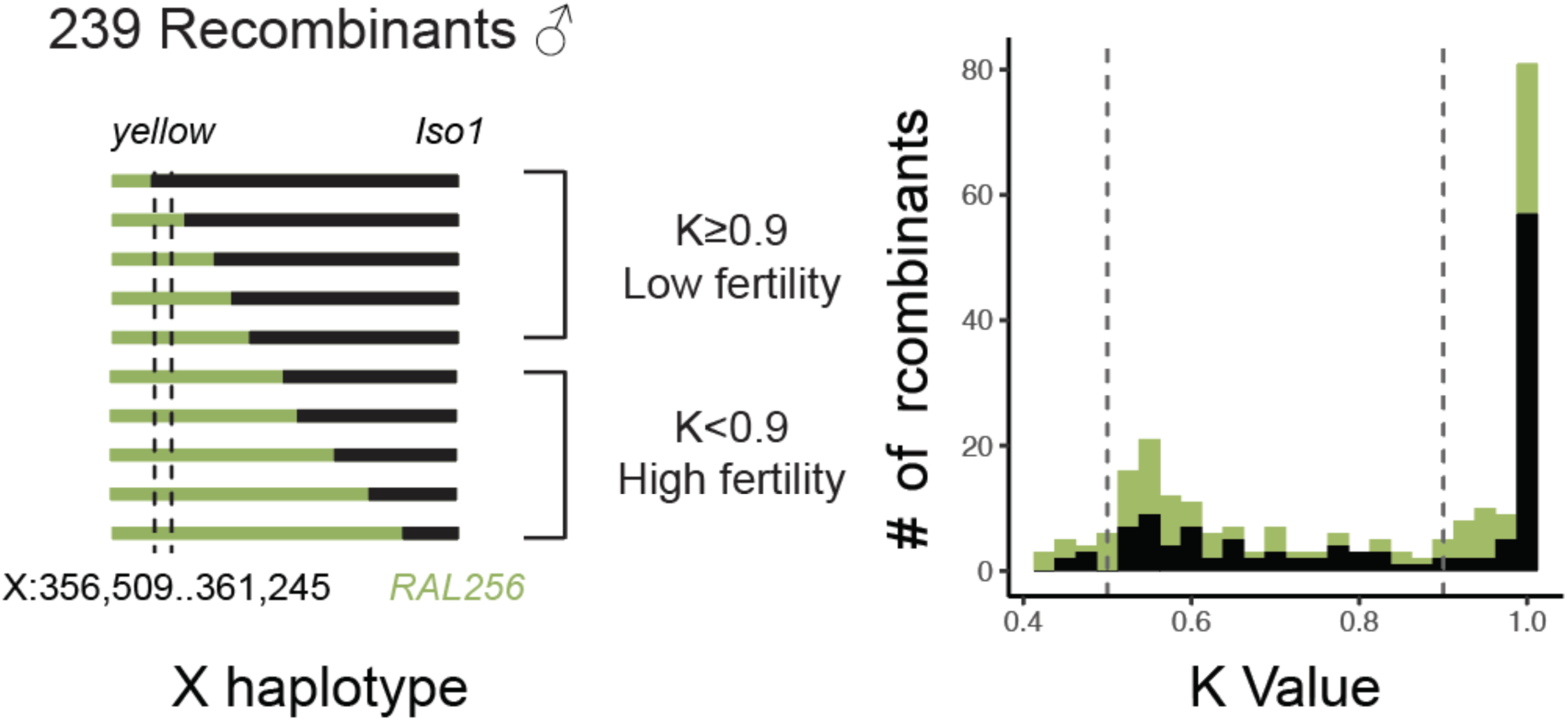
One major locus contributing to the suppressing effect of *Su(SD)X* is not linked with the visible marker *yellow*. We generated 239 recombinants between the *Su(SD)X* from RAL-256 and *yellow* chromosomes. We measured *k* values of these recombinants by counting their offspring. We also genotyped these recombinants to map suppressors. The recombining chromosomes containing *yellow* from the *Su(SD)X* chromosome are marked in green, and those containing *yellow* from the *Iso-1* are marked in black.

